# Integrative microbiome- and metatranscriptome-based analyses reveal diagnostic biomarkers for peri-implantitis

**DOI:** 10.1101/2025.06.24.661241

**Authors:** Amruta A. Joshi, Szymon P. Szafrański, Matthias Steglich, Ines Yang, Taoran Qu, Paula Schaefer-Dreyer, Xing Xiao, Wiebke Behrens, Jasmin Grischke, Susanne Häussler, Meike Stiesch

**Author notes:** correspondence to: Prof. Dr. Meike Stiesch, Department of Prosthetic Dentistry and Biomedical Materials Science, Hannover Medical School Carl-Neuberg-Str.1 30625 Hannover, Germany. Amruta A. Joshi and Szymon P. Szafrański contributed equally.

## Abstract

Peri-implantitis is a severe biofilm-associated infection of the tissues around dental implants that increases the risk of implant failure. The prognosis of peri-implantitis treatment is compromised by the resistance of well-organized and mature bacterial biofilms. Thus, early diagnosis of a pathogenic biofilm would enable treatment at a prognostically favourable stage. However, relatively little is known about how the microbial constitution of the biofilm changes during disease development. The aim of this cross-sectional study was therefore to identify peri-implant taxonomic and functional biomarkers that reliably indicate peri-implantitis using paired data from full length 16S rRNA gene amplicon sequencing (full-16S) and metatranscriptomics (RNAseq). Disease signatures were identified using 24 healthy and 24 peri-implantitis-associated biofilm samples from 32 patients. The taxonomic measurements were validated with 68 additional full-16S samples from another 40 patients. Both full-16S and RNAseq revealed significant differences between healthy and peri-implantitis samples, with respect to both the microbiome and functional profiles. A shift from aerotolerant Gram-positive bacteria to anaerobic Gram-negative bacteria was observed in peri-implantitis. Distinct metabolic pathways were expressed in healthy and peri-implantitis samples. Our results, based on paired taxonomic and functional profiles, provide for the first time important insights into the complex peri-implant biofilm ecology related to amino acid metabolism. Integrating taxonomic and functional information improved the predictive ability (AUC = 0.85) of the machine learning models and revealed diagnostic biomarkers with large effect sizes (Cohen’s d > 0.8). Primary biomarkers included health-associated *Streptococcus*, *Rothia* species and enzymes associated with peri-implantitis (urocanate hydratase, tripeptide aminopeptidase, NADH:ubiquinone reductase, phosphoenolpyruvate carboxykinase and polyribonucleotide nucleotidyltransferase - mostly expressed by Fusobacteriia and Bacteroidia). Thus, biofilm profiling at these two molecular levels reveals highly predictive disease biomarkers and provide the basis for developing early diagnostics and individualized therapy approaches for peri-implant diseases.

## Introduction

Peri-implantitis is a biofilm-associated disease that is characterized by inflammation of the peri-implant mucosa and subsequent progressive loss of the supporting bone surrounding dental implants^1,2^. It is a highly prevalent disease, with a reported rate of 22-43% across various populations within 5-10 years of implantation^3,4^. Peri-implantitis is primarily caused by an imbalance within the microbial community and the ensuing host immune response^5,6,7^. The diagnosis of peri-implantitis is founded on the presence of clinical signs and symptoms, such as bleeding on probing and increased probing depths as well as the presence of radiographic bone loss surrounding the implants^1^. However, these parameters may not reflect the biological variability across various patients and furthermore are only discernible after the onset of irreversible tissue damage^8,9^. The overall prognosis of peri-implantitis treatment at this late stage is compromised by the resistance of pathogenic biofilms to antibiotics and the lack of tissue regeneration^8,10^. It is, therefore, crucial to develop biomarkers that allow the detection of microbial dysbiosis within the biofilms early in the course of the disease, prior to the establishment of clinical and radiological changes. This would improve the early diagnosis and treatment of peri-implant diseases, when the prognosis is still favorable.

With the recent advances in next generation sequencing technologies, molecular tools could provide a promising approach for microbiome-based diagnostics^11,12,13^. So far, efforts have been focused on identifying the bacterial members of the peri-implant microbiome, and in establishing etiological links between various species and peri-implant diseases. Studies using short-read 16S rRNA sequencing and metagenomics have shown that peri-implant sites are distinct ecological niches characterized by specific bacterial communities. The transition from healthy to diseased states is accomplished by shifts in community composition rather than by the presence of specific pathogenic taxa^14,15,16,17^. However, these studies usually fail to achieve high resolution of taxonomic diversity (except metagenomics) or to achieve insights into the overall functional activities within the biofilms, as can be provided by full-length 16S rRNA gene amplicon sequencing (full-16S)^18^ and metatranscriptomics (RNAseq)^19^, respectively. This may especially be true for complex amino acid metabolism within peri-implant biofilms. This is poorly understood at the systemic level, even though it has been shown to be critical for pathogenesis and peri-implant tissue damage, as it influences the growth and survival of biofilms^20,21^.

In the search for early disease biomarkers, analyzing metatranscriptomic signatures within the complex microbiome may provide a promising approach for assessing the risk of disease development across various medical disciplines^22,23^. Furthermore, it facilitates a system-level understanding of the physiology and ecology of biofilms, as is essential for interpreting biomarkers and their interactions. However, metatranscriptomic studies are technically demanding as there may be problems with low sample biomass, RNA instability, and complex bioinformatic workflows^24,25^. Thus, only a few studies with small patient cohorts have successfully explored the microbial metatranscriptome around natural teeth^26,27,23^. There have been even fewer studies with dental implants^28,29^.

Our investigation aimed to identify microbial expression patterns and functional pathways specific for peri-implantitis by integrating high-resolution full-16S with metatranscriptomics. In the present study, we characterized the paired peri-implant taxonomic profiles and metatranscriptomes from 24 healthy and 24 peri-implantitis-associated biofilms in 32 patients. Additionally, 34 healthy and 34 peri-implantitis-associated taxonomic profiles from another 40 patients were used for validation. Our molecular analysis was empowered by a tailored genomic reference database encompassing approximately 10,000 genomes or metagenome assembled genomes. We observed profound shifts in the composition of biofilms in peri-implantitis and linked these to metabolic pathways that were differentially expressed in the two groups. System-level analysis revealed the physiology and ecology of the biofilms related to amino acids, including a network of potential microbial interdependencies. By using a machine learning approach and combining taxonomic and functional data for the first time, we identified diagnostic biomarkers for peri-implantitis with high predictive accuracy.

## Results

Our training dataset comprised taxonomic and functional data from a total of 32 participants (mean age = 67 years, 62.5% females), which are a part of the ‘SIIRI Peri-Implant Biofilm Cohort’, presenting with 24 healthy and 24 peri-implantitis-associated biofilm profiles (**Fig. 1**, **Supplementary Fig. 1**). Baseline demographics and clinical information revealed significant differences (*P*<0.0001) between healthy and peri-implantitis groups for relevant clinical measurements, including bleeding on probing, probing depth, gingival index, plaque index and suppuration (**Extended data A**). Furthermore, 68 biofilm samples (34 from each diagnosis group) from an additional cohort of 40 patients (mean age = 65 years, 52.5% females) were analyzed as a validation cohort for full-16S taxonomic biomarkers (**Extended data B**). Full-16S of the training dataset generated a total of 1,334,162 reads, with an average of 13,898 reads per sample. The identified 653 species-level taxa represented 12 phyla, 21 classes, 34 orders, 52 families and 94 genera. The species-level taxa (n = 399) that reached a relative abundance of at least 0.1% in at least one sample were considered for down-stream analysis. RNAseq using the Illumina platform produced a total of 1.5 billion reads over 48 samples, with a mean of 35.4 million reads per sample. The resulting transcripts could be associated with 1853 enzyme functions (Enzyme Commission numbers, ECs). 226 level 4 ECs reached a relative abundance of at least 0.1% in a minimum of one sample. Full-16S of the test dataset generated a total of 1,088,647 reads with an average of 16,009 reads per sample.

**Fig. 1.**
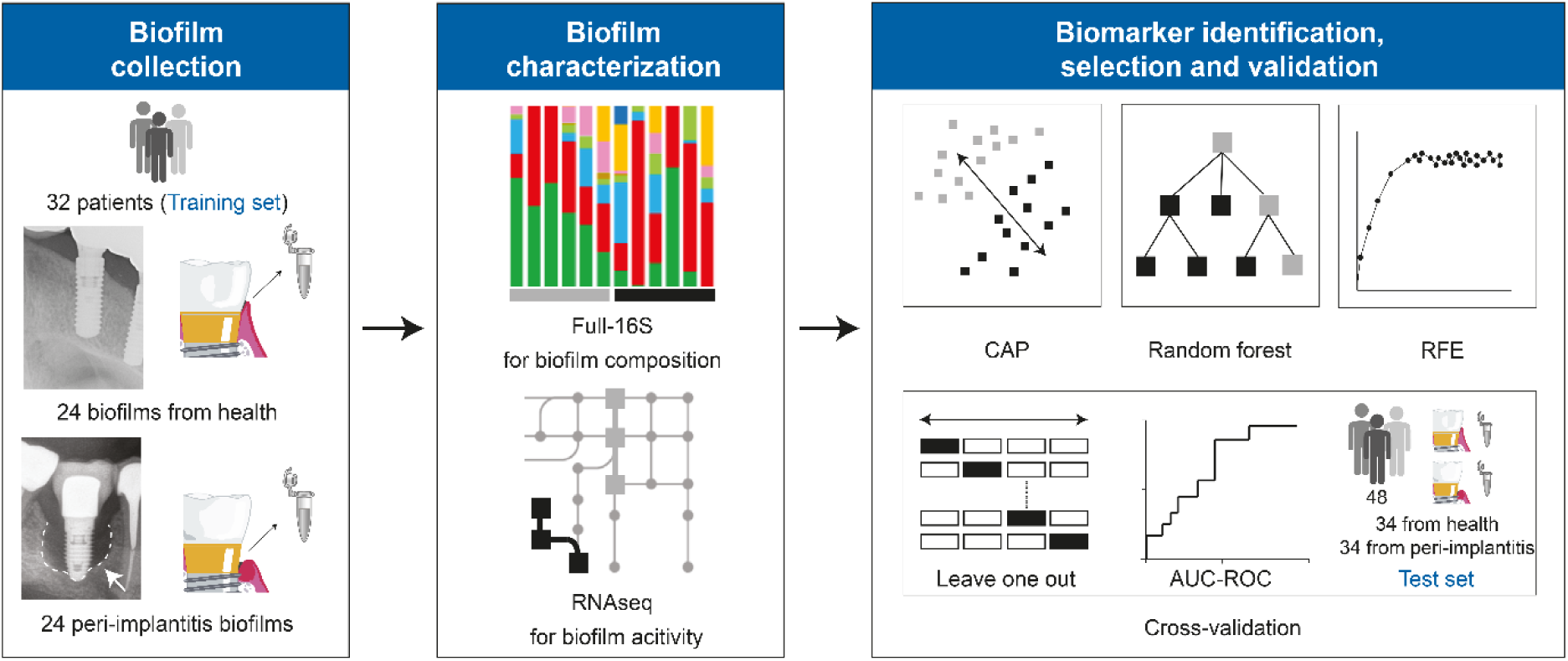
Graphical overview of the experimental design. CAP, canonical analysis of principal co-ordinates; RFE, recursive feature elimination; AUC-ROC, Area Under the Receiver Operating Characteristic Curve. Implant icon – modified from Zheng, H. et al. *Sci Rep.* **5,** 2015.

### Microbial class level abundances and functional activities were highly consistent, with the exception of Fusobacteriia and Saccharibacteria

Communities from healthy sites were dominated by members of the Bacilli class, while peri-implantitis was characterized by the dominance of Bacteroidia and Fusobacteriia - with only a few exceptions (**Fig. 2a** – **2c**) (**Extended data E** and **G**). Other classes of strictly anaerobic microorganisms (*i.e.*, Clostridia and Deltaproteobacteria) were associated with peri-implantitis. The composition of biofilms at the RNA level generally resembled the full-16S profiles, with significant variation observed between health and peri-implantitis (PERMANOVA *p* < 0.001 for full-16S and *p* < 0.05 for RNAseq) (**Fig. 2a** – **2f**). Relative abundances and relative activities correlated strongly for all classes (r > 0.75), except for Fusobacteriia (r = 0.54) and Saccharibacteria (r = −0.07) (**Fig. 2c**). The activities (RNA-level) were higher than their respective full-16S abundances for Fusobacteriia in both healthy samples and in peri-implantitis. In contrast, higher full-16S abundances were observed for Saccharibacteria than their respective activities (**Fig. 2a** – **2c**).

**Fig. 2.**
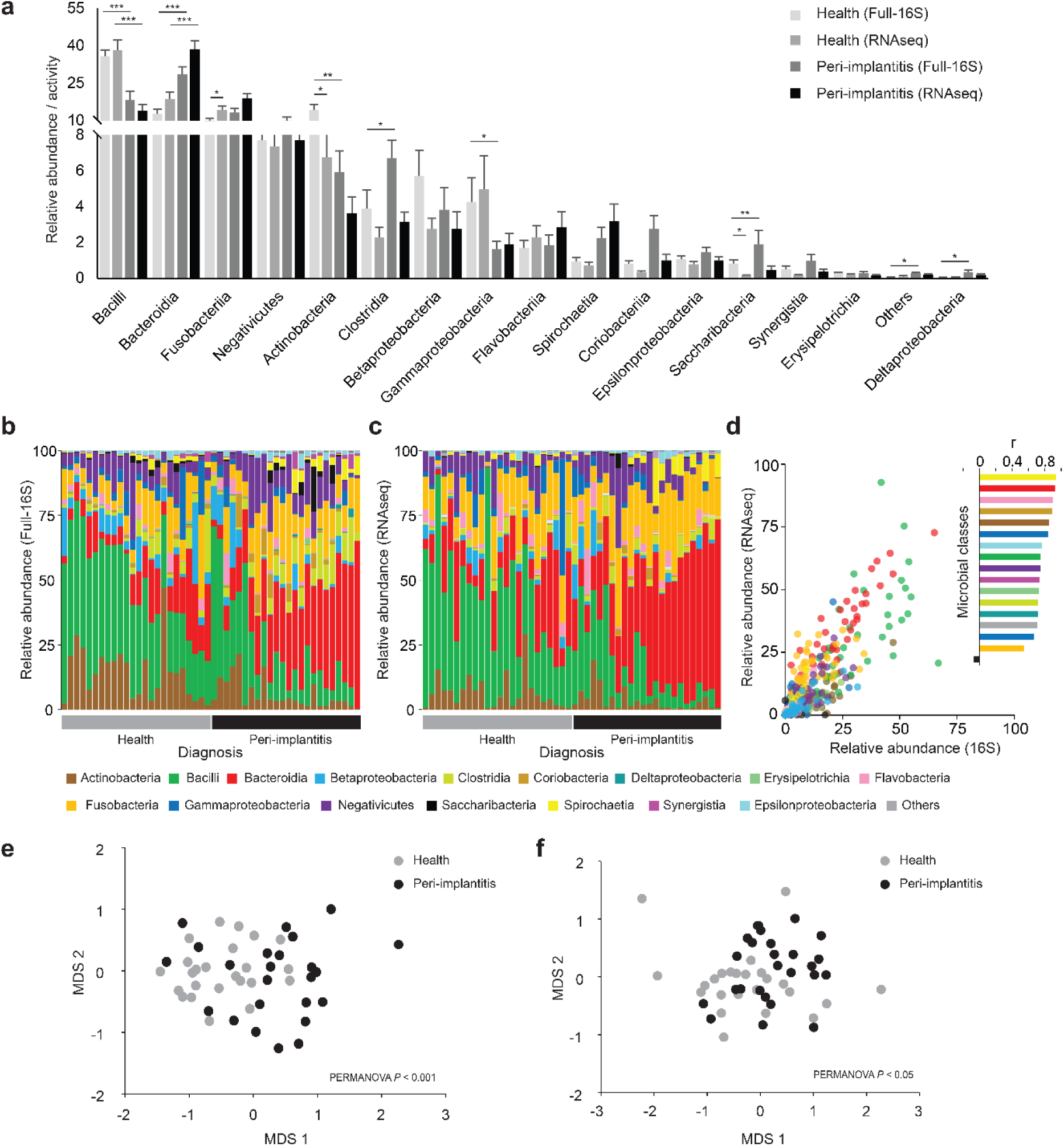
Microbial class level community composition shows distinct community abundances in healthy and peri-implantitis samples. **a** Relative abundance of taxonomic classes averaged for healthy and peri-implantitis groups in full-16S and RNAseq. The error bars show the standard errors of the means. **b** Full-16S-based bacterial class level relative abundance in each sample. **c** RNAseq-based transcriptionally active class level relative abundance in each sample. **d** Dot plots showing the correlation between the relative abundance and relative activity within biofilms with exact correlation values plotted for each class on a barplot. nMDS ordination plots based on Bray-Curtis dissimilarity matrices for class level taxa in **e** Full-16S and **f** RNAseq.

### Distinct species clusters were observed in healthy and peri-implantitis samples

**Fig. 3a** and **Supplementary Fig. 2** also show a significant shift in community composition from health to disease. In the full-16S training dataset, this was the case at both the species and genus levels (PERMANOVA, *p* < 0.0001 for both levels), as well as in the test dataset of 68 additional samples (*p* < 0.001). Aerotolerant Gram-positive species, generally regarded as benign or even beneficial in peri-implant locations (such as *Streptococcus*, *Rothia* and *Granulicatella*) dominated in health. Conversely, a high fraction of established or potential oral pathogens, including *Prevotella*, *Porphyromonas*, *Treponema*, and *Fusobacteria*, were associated with peri-implantitis. (**Fig. 3c**, **Supplementary Fig. 2**, **Extended data C and D**). **Fig. 3b** and **Supplementary Fig. 2** show results for the canonical analysis of principal co-ordinates (CAP) ordination separating the diagnosis groups at species and genus levels for the training dataset (CAP1 δ2 = 0.75 and 0.71 for species and genus, respectively). The CAP model trained on the training set successfully predicted the diagnosis of the validation set samples. Comparable accuracies were obtained for training and validation (n = 68) datasets (**Fig. 3d**). Species that showed high correlation with the CAP axis formed clusters (**Fig. 3e**). The observed species clusters also showed distinct associations to either healthy or peri-implantitis samples. In addition to the established pathogens, we observed emerging pathogens (*e.g.*, *Dialister invisus*, *Pseudoramibacter alactolyticus*, and *Fretibacterium*, *Megasphaera*, *Saccharibacteria*, *Alloprevotella* and *Selenomonas* species) in peri-implantitis (**Extended data C**).

**Fig. 3.**
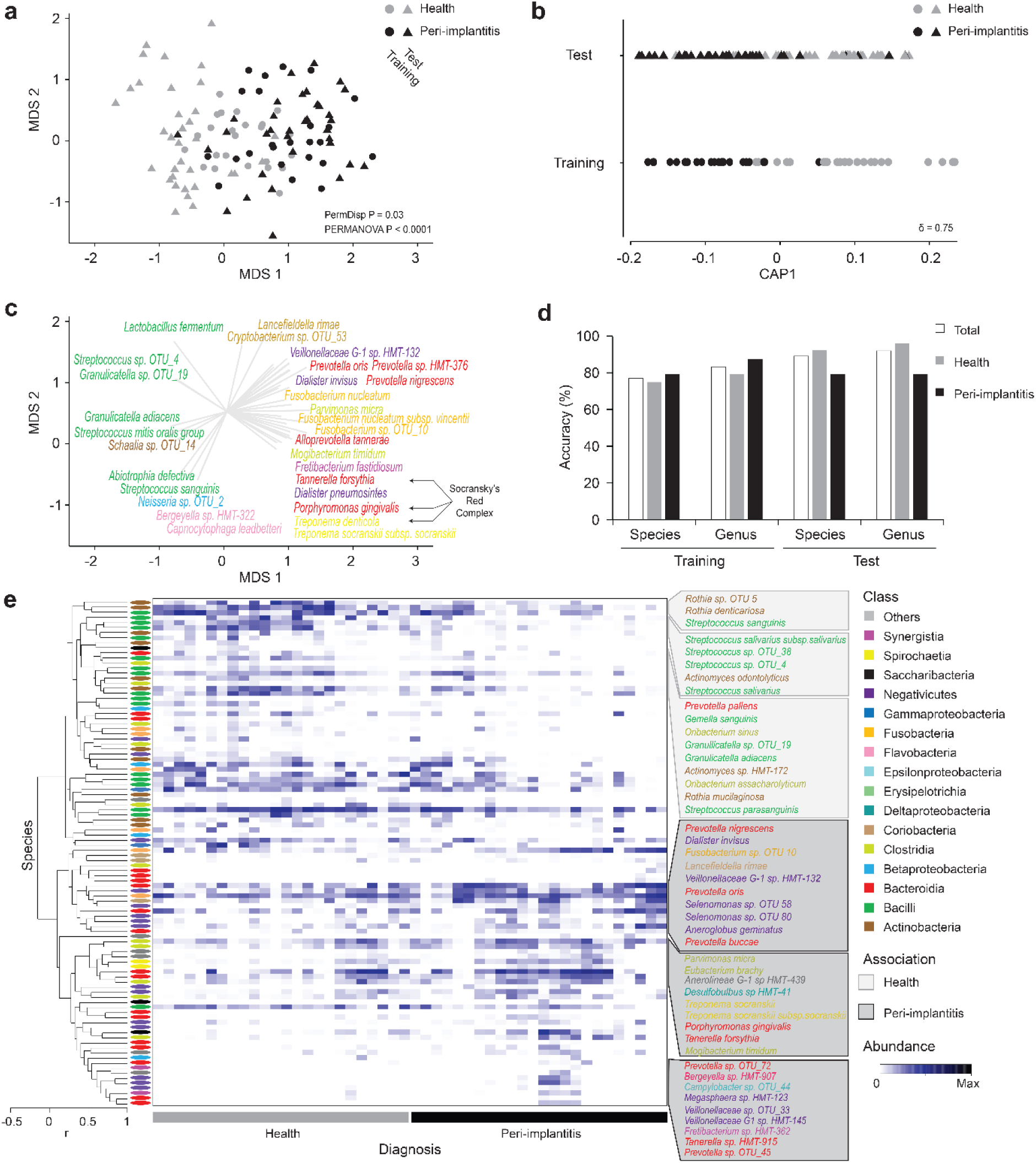
Distinct separation between diagnosis groups based on full-16S species-level taxa. **a** nMDS plots based on Bray-Curtis dissimilarity matrices of Full-16S data, aggregated to species level features of the training and validation set samples. **b** Canonical analysis of principal co-ordinates (CAP) constrained ordination plot based on species data illustrating the CAP axis that maximizes the separation of the diagnosis groups. **c** Vector overlays on nMDS plot shows the species with highest correlation to the CAP1 axis. Species are coloured by bacterial classes. **d** Results of leave one out cross-validation of CAP analysis showing proportions of correct allocations of observations to diagnosis groups across species and genus level taxonomic features in training and test datasets. **e** Heatmap showing the standardized signal for selected 16S taxonomic biomarkers with the highest correlation to CAP1 axis. Clustering of species level taxa is based on Spearman Rank correlation matrix.

### Diagnosis-specific differences in functional activities revealed differentially expressed metabolic pathways

To gain a better understanding of the prominent microbial activities, we visualized differentially expressed enzyme classes (ECs) on the metabolic pathways map provided by the Kyoto Encyclopedia of Genes and Genomes (KEGG) (**Fig. 4a**). Unconstrained ordination showed distinct differences between diagnosis groups (PERMANOVA, *p* <0.0001) as assessed by taxon-independent RNAseq reads grouped to ECs (**Fig. 4b**). The CAP model, built on functional data, effectively separated diagnosis groups (CAP1 δ2 = 0.65) (**Fig. 4c, Extended data F**). The volcano plot (**Fig. 5a**) illustrates 277 and 221 differentially expressed ECs (*p* < 0.001; CAP |r| > 0.4) in healthy and peri-implantitis samples, respectively. The BRITE classification of these ECs revealed the most abundant differentially expressed pathways across diagnosis. Important metabolic pathways related to amino acids (histidine, alanine, aspartate, glutamate, cysteine, methionine, tryptophan, glutathione, phenylalanine, tyrosine, lysine, valine, leucine, isoleucine and butanoate) were the most abundant differentially expressed pathways across diagnosis groups. Furthermore we detected many other metabolic pathways that were either associated with health (pyruvate metabolism, galactose metabolism, sulphur metabolism, starch and sucrose metabolism) or peri-implantitis (lipopolysaccharide biosynthesis, amino-acyl t-RNA biosynthesis, O-antigen nucleotide sugar biosynthesis, carbon fixation and one carbon pool by folate) (**Fig. 5b**).

**Fig. 4.**
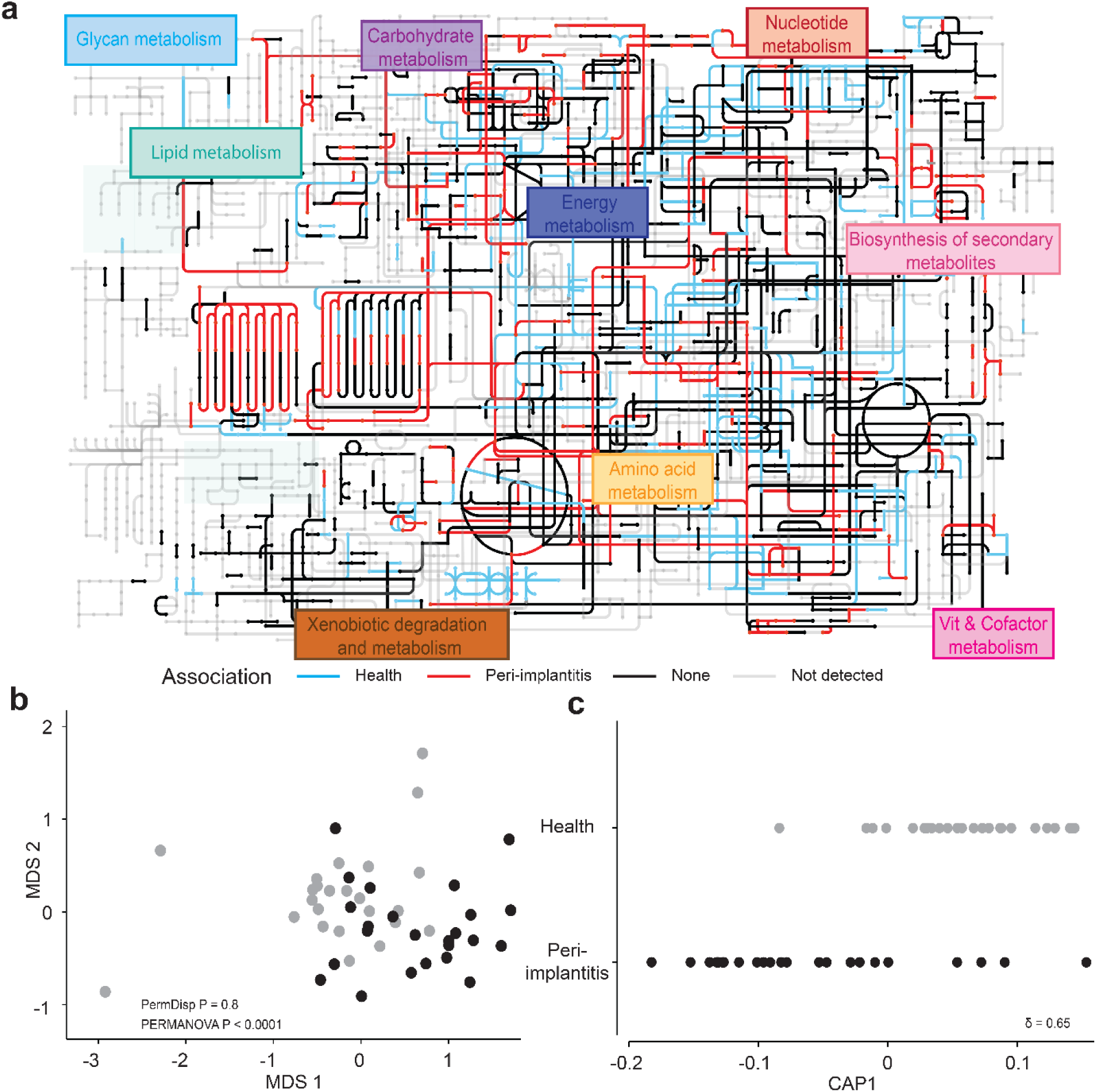
Distinct metabolic pathways were expressed in healthy and peri-implantitis samples. **a** Global overview of metabolic pathways generated using KEGG annotation of the RNA based ECs and their expression in health and peri-implantitis. **b** nMDS plots based on Bray Curtis dissimilarity matrices of log (x+1) transformed RNA data, aggregated to EC4 level **c** CAP constrained ordination plots of EC4 level data illustrating the CAP axis that maximizes the separation of the diagnosis groups.

**Fig. 5.**
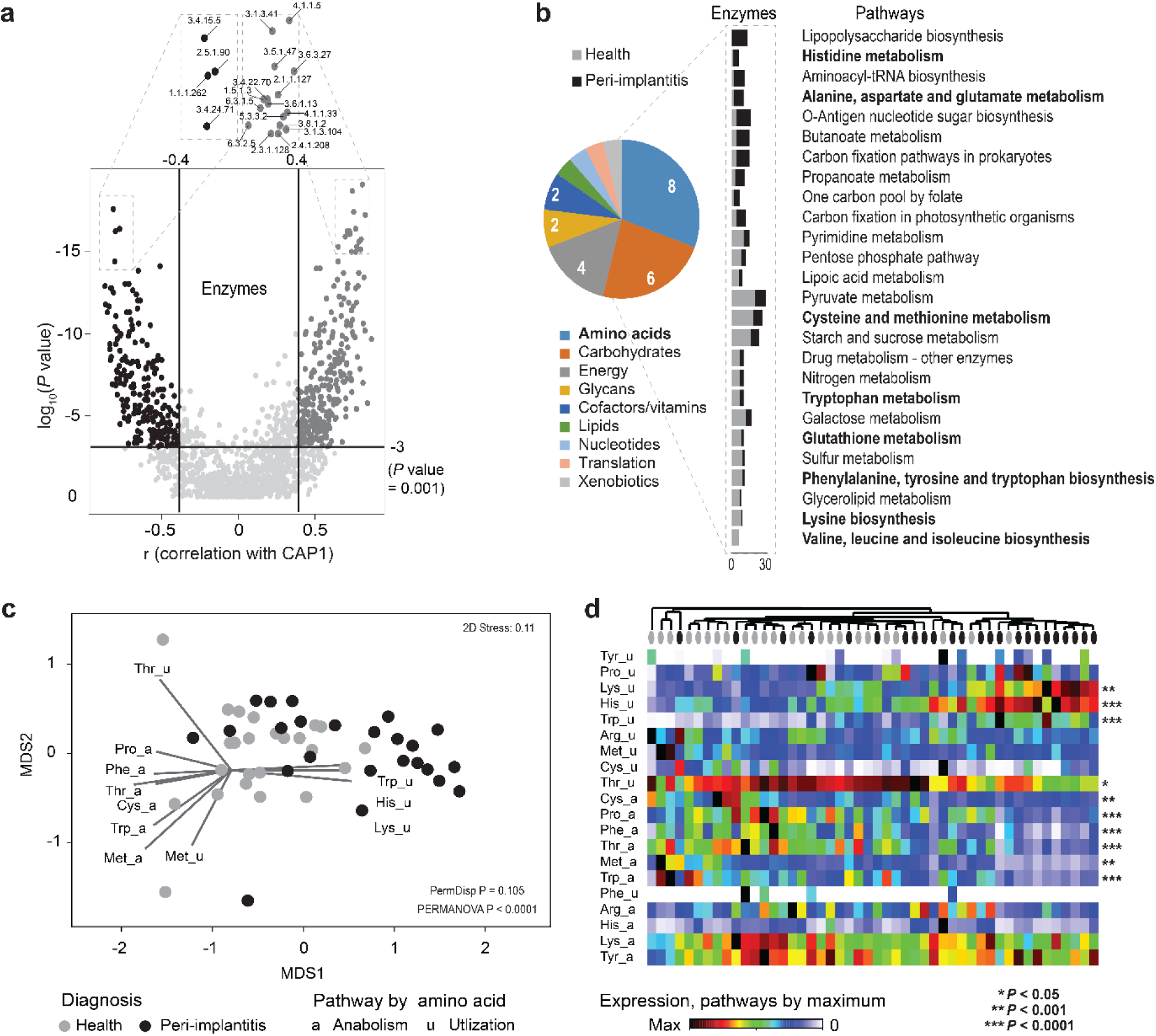
Amino acid metabolic pathways as the most differentially expressed pathways between diagnosis groups. **a** Volcano plot of differentially expressed ECs in health and peri-implantitis based on the LDA correlation cut-off ± 0.4 and *p* value ≤ 0.001. **b** Specific metabolic pathways identified using the BRITE classification of ECs, revealed amino acid metabolic pathways as the most abundant differentially expressed pathways between the diagnosis groups. **c** nMDS ordination plot based on Bray-Curtis matrix of normalized EC counts associated with different amino acid anabolic and utilization pathways. (PERMANOVA *p* value ≤ 0.0001) **d** Heat map showing the abundances of RNA-based reads grouped to amino acid metabolic pathways. Samples clustered based on the Bray-Curtis similarities. *p* values are obtained by Mann-Whitney U test with FDR correction, calculated for two diagnosis groups.

### Peri-implantitis is associated with increased utilization of histidine, lysine and tryptophan

To further explore the fifteen amino acid related pathways that were differentially expressed in healthy and peri-implantitis groups, we created a curated amino acid metabolic pathway dataset (**Supplementary Fig. 4**, **Methods, Extended data H**) The nMDS plot based on this dataset showed considerable separation between diagnosis groups with PERMANOVA *p* < 0.0001 (**Fig. 5c**). High correlation of histidine, lysine and tryptophan utilization was observed with peri-implantitis samples while anabolic pathways were associated with healthy samples. Additionally, another axis was uncovered that represented complementary anabolism and utilization of methionine. Clustering showed distinct grouping of healthy and peri-implantitis samples, while the accompanied heatmap showed the major gradients in activities and many unique pathway profiles (**Fig. 5d**). Pathways with high correlation to the CAP axis also showed consistent diagnosis-specific signals. In contrast, arginine and histidine anabolism and threonine utilization were more uniformly expressed throughout the samples, irrespective of the diagnosis. Utilization of aromatic amino acids (tyrosine and phenylalanine) was represented as a unique characteristic of a few samples.

### Prevalence of potential amino acid dependencies in peri-implant biofilms indicate metabolic exploitation

Integrating the amino acid metabolic pathways with taxa that actively transcribed the relevant genes led to a systemic understanding of the amino acid physiology and ecology within peri-implant biofilms and highlights the network of key metabolites (**Fig. 6a**). The high resolution view of the pathway for individual amino acids (**Fig. 6a**, **Supplementary Fig. 4 and 5**) illustrated consistent expressions of all pathway enzymes and of the taxonomic classes transcribing the relevant genes. Interestingly, the predicted biosynthesis and utilization of amino acids by the different bacterial species could be linked and served to visualize a sequential network that resulted in end-products with different pathogenic potentials, such as short chain fatty acids (*e.g.*, acetate, butyrate, propionate) or polyamines (such as putrescine) (**Fig. 6a**). Diverse peptidases (*i.e.*, enzymes that produce amino acids and small peptides) were expressed in biofilms and many of these enzymes were associated with peri-implantitis (**Fig. 6b**). Eight out of 10 amino acids (exception: tyrosine, methionine) showed substantial differences (log2 folds cut-off > 1.5) between anabolic/catabolic activity versus predicted total activity (**Fig. 7a**, **Supplementary Fig. 6a and 6b)**. Negativicutes, Bacilli and Actinobacteria were the predicted major amino acid producers, while Fusobacteriia, followed by Flavobacteria, were predicted to be more dependent on amino acid supply. Phenylalanine and histidine showed the highest number of connections to different bacterial classes.

**Fig. 6.**
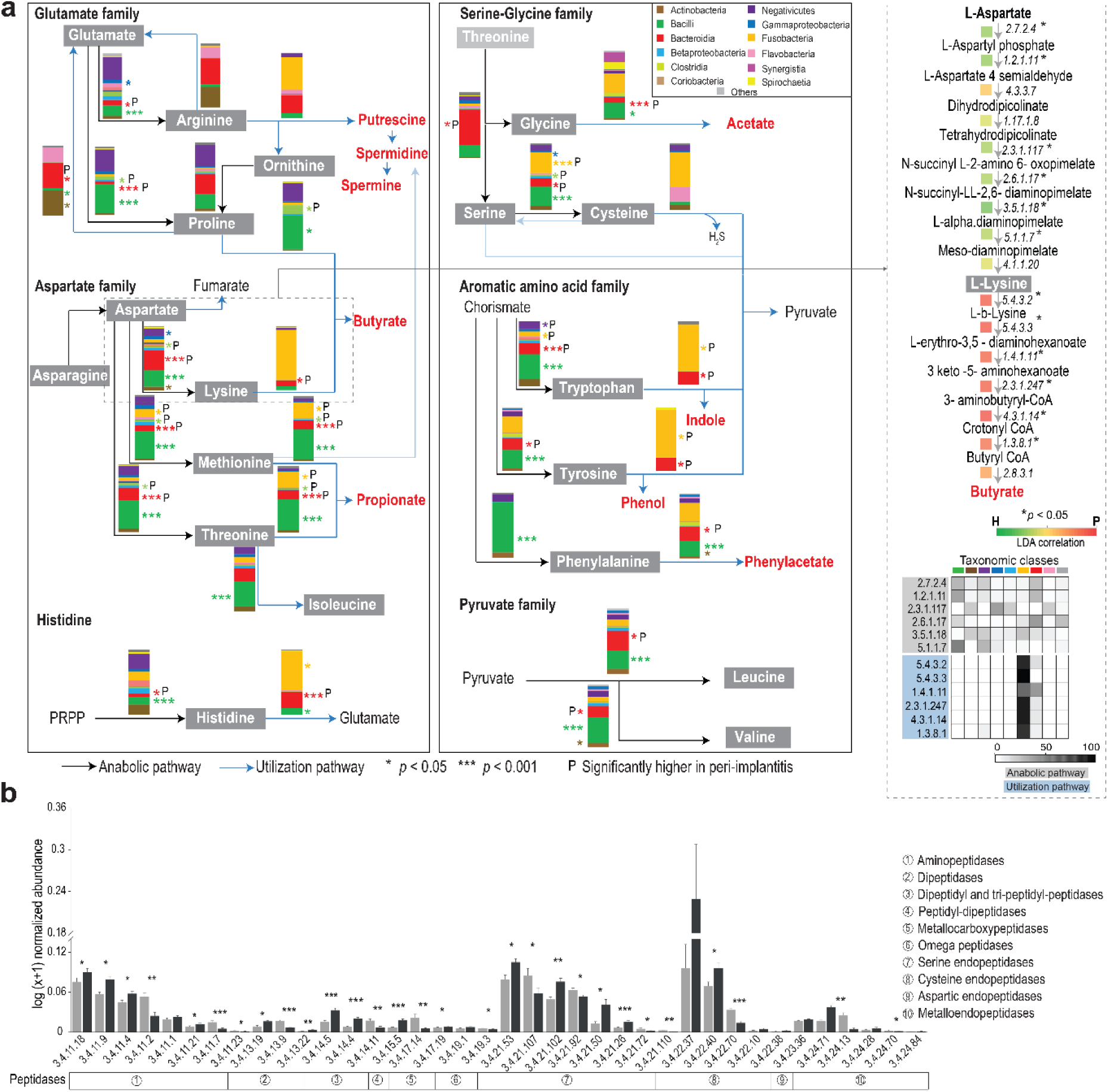
System-level perspective of amino acid ecology within peri-implamt biofilms. **a** Proposed bacterial metabolic pathways for amino acids production and utilization based on metatranscriptomic profiles obtained in this study. Accompanied bars indicate the average relative abundances of major bacterial classes contributing to the respective amino acid activities. Lysine as a model illustrating the high-resolution view of the curated pathways based on RNA ECs. Color scale indicates the LDA correlation (CAP model based) of each enzyme to the CAP1 axis that best separates the diagnosis groups. Heat map showing the standardized signal for lysine anabolic and utilization activities within the important bacterial classes. **b** Normalized abundances of peptidases in healthy and peri-implantitis samples. *p* values are obtained by Mann-Whitney U test with FDR correction, calculated for two diagnosis groups. * *p* < 0.05, ** *p* < 0.01, *** *p* < 0.001.

**Fig. 7.**
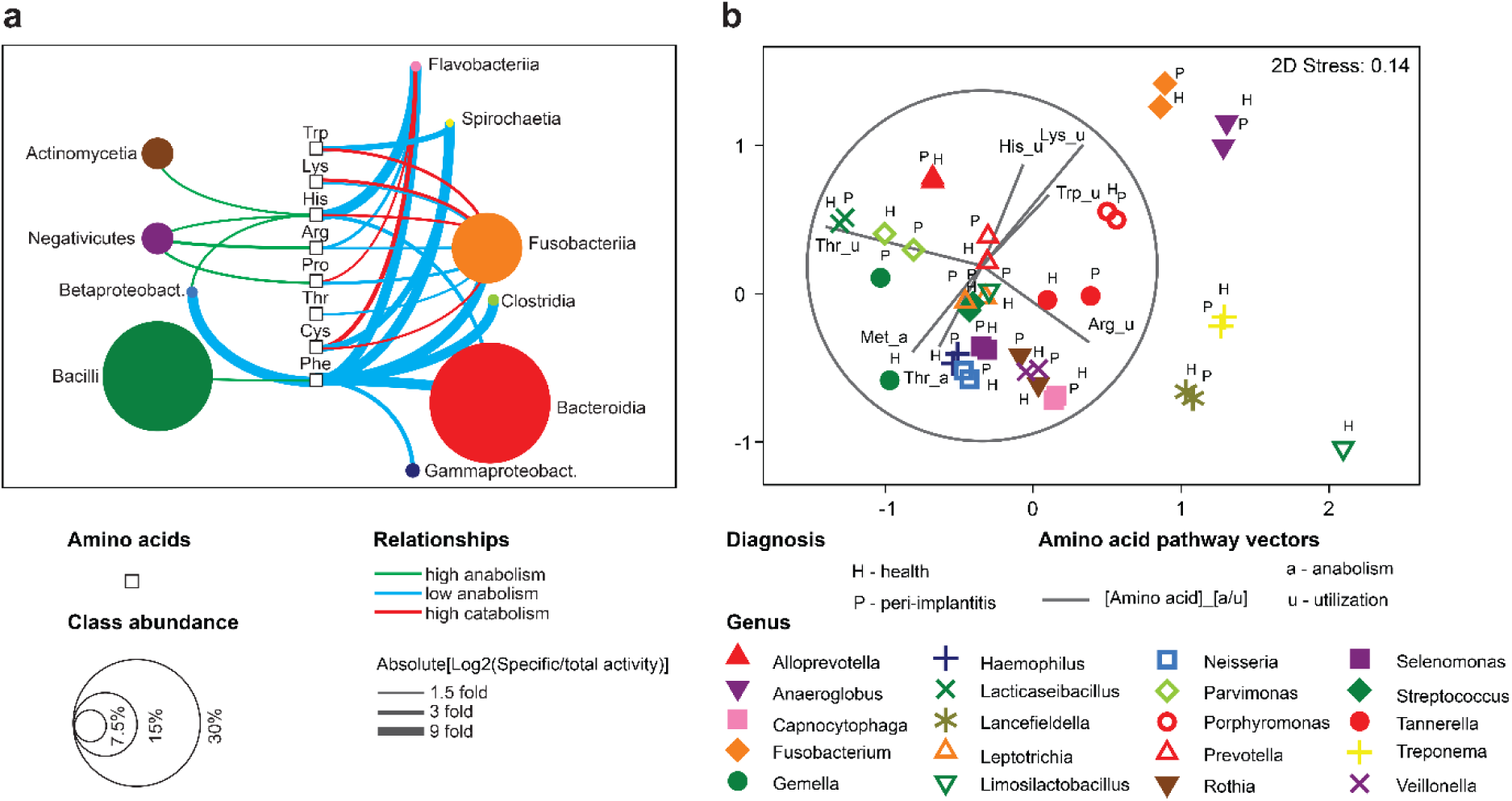
Predicted amino acid activities of major bacterial classes. **a** Levels of amino acid anabolic and catabolic activities in diverse taxa visualized as network comprised of 8 amino acids. **B** Amino acid activities of the most abundant genera in both diagnosis groups. See **Supplementary Fig. 6** for more details.

Metabolic strategies of specific taxa were studied at higher resolution, in order to capture intraclass differences and aid interpretation. Different metabolic strategies of the 20 most active genera were observed, as depicted by four major gradients **(Fig. 7b**, **Supplementary Fig. 6c)**. Peri-implantitis-associated utilization pathways of histidine, lysine and tryptophan were linked to *Porphyromonas*, *Anaeroglobus* and *Fusobacterium*. Although genera representing the same classes usually co-localized (*e.g.*, Bacilli, Bacteroidia), intraclass differences were obvious (*e.g.*, *Porphyromonas, Prevotella*, *Tannerella*), which suggests that competition is minimized via niche selection. High intraclass differences were also observed in Fusobacteriia (*Fusobacterium* vs *Leptotrichia*) and Negativicutes (*Anaeroglobus* vs *Selenomonas* and *Veillonella*). Occasionally, taxonomically unrelated taxa converged at the same utilization strategies (*e.g.*, *Lancefieldella* and *Treponema*), indicating an excess of amino acid substrate. Taxon-specific strategies at genus level were usually consistent for diagnosis groups, except *Gemella* and *Limosilactobacillus*, that showed some variation.

### Combined microbiome-metatranscriptome classifiers can accurately predict peri-implant diagnosis

We next aimed to combine DNA- and RNA-based information for a machine learning algorithm to identify potential biomarkers for peri-implantitis diagnosis. For this purpose, we created datasets comprising the most differentially expressed features (high correlation to CAP axis) across five data sets, consisting of species, genus, EC alone as well as their combinations. and used this to train a random forest (RF) prediction model. The model performance as assessed by ‘20X repeated 10-fold cross-validation’ demonstrated improved performance of the combinations compared to the individual datasets. The AUC values ranged from 0.85 for the ‘Species + EC combination’ model to 0.64 for the ‘genus level’ model. The AUC values for species (0.83), genus + EC (0.81) and EC alone (0.74) were lower than the ‘Species + EC combination’ that performed the best. (**Fig. 8a**). Most biomarker candidates were consistent across datasets (**Fig. 8b**). **Fig 8c** shows the top biomarkers from the ‘Species + EC combination’ dataset. *Streptococcus* species OTUs that matched best with *S. salivarius* and the understudied *Streptococcus* species HMT-074 (related to *S. oralis*) were identified as the important taxonomic biomarkers, along with *Rothia dentocariosa*. Polyribonucleotide nucleotidyltransferase (PNPase), Na+ transporting NADH:ubiquinone reductase (Na+NQr), phosphoenolpyruvate carboxykinase (PEPCK), tripeptide aminopeptidase and urocanate hydratase were identified as top enzyme biomarker candidates for the diagnosis. The normalized abundances of these top biomarkers plotted for health and peri-implantitis (**Fig. 8d**) showed significant differences (FDR corrected *p*< 0.05, except for tripeptide aminopeptidase), with large effect sizes of Cohen’s d statistic of > 0.8.

**Fig. 8.**
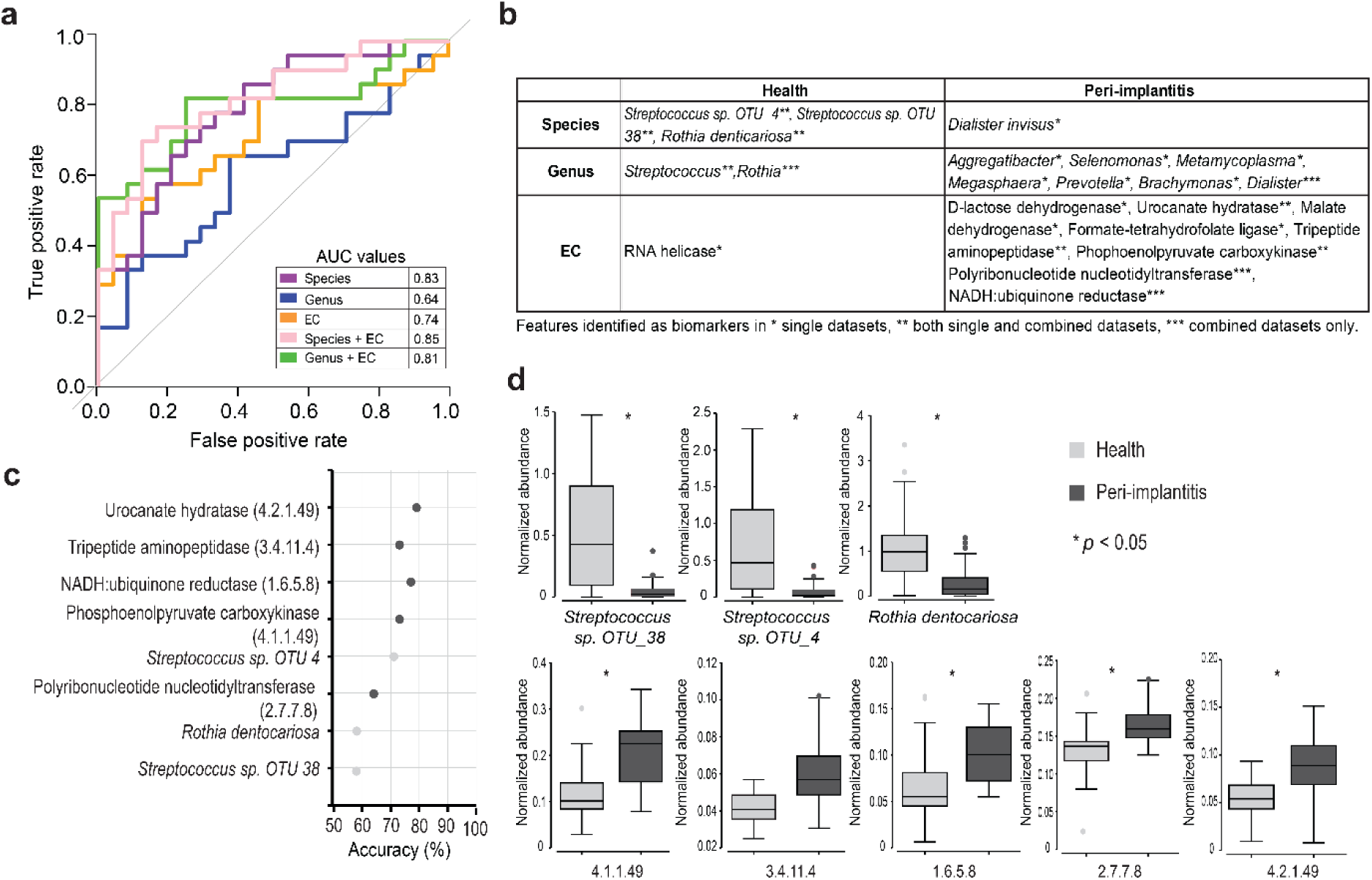
Random Forest (RF) predictive modelling results. **a** Results of AUC-ROC and LOOCV accuracies comparing RF models based on different inputs for diagnosis prediction. **b** Top biomarker candidates for health and peri-implantitis identified by different input datasets. **c** Results of RFE showing the optimum number of features from ‘Species + EC’ combination. **d** Normalized abundances of top biomarkers between diagnosis groups. * indicates *p* values obtained by Mann-Whitney U test with FDR correction, calculated for two diagnosis groups.

## Discussion

With an increasing number of patients receiving dental implants globally^30^ and the high prevalence of peri-implant disease^3,4^, there is an urgent need to identify and validate biomarkers for early diagnosis of peri-implantitis. Our high throughput study is the first to provide a paired full-16S and metatranscriptomic characterization of peri-implant biofilms and to identify an optimal combination of DNA-RNA biomarkers with strong predictive abilities for peri-implantitis diagnosis.

Overall, strong correlation was observed between full-16S and RNAseq - except for Saccharibacteria and Fusobacteriia. Both methods revealed consistent shifts in the biofilm community, from Gram-positive facultative commensals in health to expansion of Gram-negative anaerobic pathogens in peri-implantitis. This is in agreement with the results of previous studies^28,31,32,33^. However, the superior discriminatory power of full-16S conferred by circular consensus sequencing (CCS) of long amplicons enabled us to assign species taxonomy to a higher percentage of OTUs^18^. Species clusters of emerging pathogens (*e.g, Dialister invisus*, *Pseudoramibacter alactolyticus*, *Fretibacterium*) observed in a few peri-implantitis samples have also been reported in recent studies^16,17,34,35^. Understanding how these species contribute to peri-implant pathologies may aid the development of individualized treatment in the future. The lower correlation between full-16S and RNAseq for Saccharibacteria could be attributed to the underrepresentation of its genomes in the reference database. However, for Fusobacteriia, it suggests that both methods are indispensable and provide complementary information. While full-16S provides a cost-effective and reliable taxonomic description, RNAseq supports a comprehensive functional understanding of the key players and their activities within biofilms^25^.

The KEGG based functional pathways analysis revealed that there are significant differences in the transcriptional landscapes of the healthy and the peri-implantitis microbiome. In particular, our results demonstrate a large shift in the metabolism of amino acids between the diagnosis groups. This indicates that the subgingival environment of dental implants is a dynamic oral niche. In health, nutrients are mainly derived from saliva and occasionally from meals, with strong fluctuations in availability^36^. In peri-implant diseases, large amounts of nitrogenous compounds (proteins and their degradation products: peptides and amino acids) occur in the destructed tissues and provide a nutrient-rich environment for microorganisms. Certain proteolytic and peptidolytic bacteria break down these amino acids to obtain energy^37^. In addition, amino acid metabolism leads to the production of end metabolites, such as short-chain fatty acids, ammonia and polyamines. These compounds in turn contribute to pH control and further tissue destruction in peri-implant diseases^21,38,39^.

With our curated approach, we could achieve system-level understanding and prediction of the physiology and ecology of peri-implant biofilms, as related to amino acid anabolism and catabolism. Our results showed that amino acid anabolism is associated with healthy states, indicating the efforts of the biofilm members to balance the scarcity states within the biofilms^40^. In peri-implantitis, significantly higher utilization of histidine, lysine and tryptophan was observed, and this is consistent with previous metatranscriptomic studies on periodontitis^26,27^. In this study, we showed for the first time that these utilization pathways are predominantly transcribed by Fusobacteriia, followed by Bacteroidia. Furthermore, at the class and genus levels, we demonstrated that different taxa choose strikingly different strategies in their amino acid metabolism to adapt to their environment. Lack of transcriptional activities related to anabolism of certain amino acids in specific taxa implies that there are metabolic dependencies through leakage between different biofilm members^41,42^. An example of this is the low anabolism and high catabolism observed in Fusobacteriia in our data, suggesting its metabolic dependency on Negativicutes and Bacilli for amino acid supply^43^. Our results provide new insights into amino acid metabolism within peri-implant biofilms by predicting the network of taxa contributing to amino acid pathways at a transcriptional level. Such dependencies of pathogens may be exploited in the future for individualized prevention strategies by shifting the balance towards health-associated microorganisms.

Our random forest machine learning classifier identified an optimal DNA-RNA biomarker combination of eight features. The majority of studies so far have employed differential expression analyses that reveal an exhaustive list of discriminatory features^35,44,45^. As the random forest method can reduce the number of features, it may be particularly well suited for developing practical biomarker panels for clinical application. The DNA-RNA combined datasets also improved the diagnostic accuracy of predictive machine learning models. This is in accordance with gut microbiome studies that have shown that metatranscriptomic profiles, with their higher variability and sensitivity, complement DNA-based information in distinguishing between healthy and diseased states^22,46^. All three taxonomic biomarkers (species related to *S. salivarius* and *S. oralis*, and *Rothia dentocariosa)* are the constituents of the core oral microbiota^47^, and usually exhibit a competitive advantage over pathogenic species^48,49,50,51^. The identified enzyme biomarkers PNPase, PEPCK and Na+NQr are involved in the metabolic adaptation mechanisms of oral bacteria in the inflammatory peri-implant environment, such as nutrient acquisition and stress response, and may contribute to the fitness of the pathogenic biofilm in peri-implant pockets^52,53,54^. Additionally, urocanate hydratase and tripeptide aminopeptidase, which are directly involved in amino acid metabolism, may contribute to the increased virulence of pathogenic species^55,56^. The development of new diagnostics to detect these microbial enzymes in a clinical setting requires the introduction of specific enzyme-targeting methods using antibodies and chromogenic products. Microbiome diagnostic tests targeting the DNA counterparts of these enzymes (*e.g.*, PCR- or hybridization-based) could be another precise and cost-effective strategy for the development of diagnostic biomarkers.

Our study is the first to address an unmet clinical need for diagnostic biomarkers for peri-implantitis through paired full-16S and RNAseq in a sizable patient cohort and successfully identified molecular biomarkers for peri-implantitis. Both full-16S and RNAseq revealed consistent shifts in microbial community composition between health and peri-implantitis. However, certain disease-specific biomarkers were revealed at the level of taxon independent enzymes (ECs) or pathways. Furthermore, the reconstruction of the amino acid ecological interaction network using metatranscriptomics revealed its significance for biofilm fitness (development of niche specialization and metabolic co-operation) and virulence (production of SCFAs and polyamines). System level analysis is a crucial step in biomarker development to establish the mechanistic links of potential biomarkers with other components of molecular pathways and to interpret their role within complex biofilm communities.

In summary, the combination of full-16S and RNAseq increased the predictive power for diagnosing peri-implantitis. Three health-associated species and five peri-implantitis-associated ECs were identified as powerful diagnostic biomarker candidates. These results indicate that measuring microbial activity within peri-implantitis-associated biofilms provides more informative insights than taxonomic profiles alone. Our integrated multi-omics approach provides a significant step towards the development of personalized chair-side early diagnostics for peri-implantitis.

## Material and methods

### Study design and ethical statement

The present prospective, clinical, cross-sectional study was conducted at the Department of Prosthetic Dentistry and Biomedical Materials Science of Hannover Medical School, Germany. The study was part of the ‘SIIRI Peri-Implant Biofilm Cohort’. SIIRI (Safety Integrated and Infection Reactive Implants) is an interdisciplinary consortium that will recruit hundreds of individuals and monitor their peri-implant microbiome for a period of up to 12 years to study the development of peri-implant diseases and to develop strategies for their early detection and prevention. The study was designed and conducted in accordance with ‘The Strengthening the Reporting of Observational studies in Epidemiology (STROBE)’ guidelines for conducting observational studies. The study protocol was in accordance with the Helsinki Declaration, as revised in 2013 and was approved by the institutional ethics committee of the Hannover Medical School (no. 9477). The written informed consent was obtained from all the patients diagnosed with either peri-implant health or peri-implantitis according to the case definitions given by the World Workshop on the Classification of Periodontal and Peri-implant Diseases and Conditions, 2017.^5^ Peri-implant health was defined as absence of bleeding on slight probing, pocket depth of ≤ 5mm and the absence of further additional bone loss following initial healing as seen on radiographs. Peri-implantitis was defined as the presence of bleeding on probing, peri-implant probing depths ≥ 6mm and radiographic evidence of bone loss around implants ≥3 mm. Exclusion criteria were presence of systemic diseases, use of antibiotics within last 3 months, and use of immunosuppressant drugs within last 3 months.

### Clinical examination

A total of 79 implants from 32 participants were included in this study. Additional 48 implants from 40 participants were included for validation of full-16S results. Validation cohort is also a part of ‘SIIRI Peri-Implant Biofilm Cohort’ and included 10 implants from longitudinal analysis (Dieckow S *et al.* - submitted to npj Biofilms and Microbiomes). Detailed medical and dental histories were recorded for all individuals by an experienced clinician who also performed the chair-side examination to record the clinical implant parameters by periodontal probing (UNC-15, HuFriedy Mfg. Co. LLC, Chicago, USA). The primary meta-data included bleeding on probing (BOP), peri-implant probing depth (PD), suppuration, gingival index (GI), plaque index (PI) and radiographic bone loss. The following secondary parameters were also recorded: i. age and gender, ii. presence of periodontitis, iii. history of periodontitis, iv. smoking status, v. residual teeth, vi. type of superstructure (fixed or removable), vii. implant age (in function). Descriptive statistics and clinical characteristics of the participants from main and validation cohort are summarized in the extended data in the online version.

### Peri-implant biofilm sample collection and processing

Prior to sample collection, all the patient information was anonymized and the participants were assigned unique patient IDs and sample IDs. The peri-implant submucosal biofilm sampling was performed according to our previous study^57^. In brief, sterile paper points (ISO 35/2.0, VDW GmbH, München, Germany) were inserted into the peri-implant sulcus or pocket at six different surfaces (mesio-buccal, mesial, mesio-lingual, disto-lingual, distal and disto-buccal) for 30 sec each, after isolating and drying the cervical region using cotton pellets. Additionally, subgingival plaque was collected at these sites using periodontal curette (HuFriedy Mfg. Co. LLC, Chicago, USA) and was transferred to the paper points before pooling them for each implant and incubating in RNAprotect (Qiagen, Hilden, Germany) for 5 min at room temperature. The samples were then immediately stored at −80 °C until further processing. RNA isolation from the biofilm samples was performed as per previous protocol^26^. The protocol was modified to include co-isolation of DNA from the same samples for full-length 16S rRNA gene amplicon sequencing. Accordingly, chemicals were obtained from Sigma-Aldrich (Sigma-Aldrich, Taufkirchen, Germany) and the kits were used according to the manufacturer’s instructions, if not stated otherwise. All glassware and other instruments were RNase-decontaminated using RNase ZAP solution (Ambion, Austin, TX, USA). The paper points were thawed and shredded with sterile scissors. The fragmented paper points were incubated in lysis buffer containing 10 mM Tris, 1 mM EDTA, pH 8.0, 2.5 mg/ml lysozyme and 50 U/ml mutanolysin at 25 °C for 1.5 h on a shaking incubator at 350 r.p.m. A total 700 μl of fresh buffer RLT (Qiagen) containing 1% (v/v) β-mercaptoethanol was added and vortexed for 10 sec. Samples (including the fragmented paper points) were placed on a QIAshredder Mini Spin column (Qiagen) and centrifuged for 1 min at 11,000 r.p.m. The flow-through containing the bacterial cells was mixed with 150 mg acid-washed and autoclaved glass beads (diameter 106 μm). Samples were vortexed 10 times for 30 sec at full speed with at least 1-min intervals on ice in between vortexing. The samples were then centrifuged for 1 min at maximal speed. Total RNA was isolated from the supernatants using the RNeasy Mini Kit (Qiagen). DNA was removed by column digestion and by DNase digestion in the eluate using the RNeasy cleanup procedure.

Once the eluate for RNA extraction had been separated, DNA isolation was performed by placing the RNAeasy minispin columns in new tubes and incubated with 30 µl of 8 mM NaOH at 55 °C for 10 min. Eluate was collected by centrifugation and 3.03 µl 0.1M HEPES was added to it and stored at −80 °C until further use.

Additional pre-processing steps were performed for samples undergoing RNAseq. Total RNA samples with low quantity of RNA which belonged to the same patient with the same clinical diagnosis and comparable clinical parameters were pooled at the stage of total RNA to meet inclusion criteria for downstream analyses. This resulted in total of 24 RNA samples from 55 healthy samples. These 24 samples were further processed similar to other unpooled samples. The DNA counterparts of these 55 healthy samples were processed separately according to the protocol thus yielding 55 DNA samples from healthy samples. The generated full-16S reads from these samples were averaged during the downstream analysis to maintain consistency between RNA and DNA features (24 healthy and 24 peri-implantitis biofilm profiles).

### Analysis of microbial composition using full length 16S rRNA gene amplicon sequencing

A previously published protocol was followed for examining the composition of the biofilms using PCR amplification and PacBio Sequencing^58^. In brief, amplification of the full-length 16S rRNA gene amplicons extracted from each sample was performed using 27F: AGRGTTYGATYMTGGCTCAG and 1492R: RGYTACCTTGTTACGACTT universal primer set. The KAPA PCR mix was used to perform 23-27 cycles of PCR amplification, with denaturing at 95 °C for 30 sec, annealing at 55 °C for 30 sec and extension at 72 °C for 90 sec. This was then followed by final synthesis for 10 min at 72 °C. Post-amplification quality control was performed using Invitrogen Qubit dsDNA and BR Assay Kit (Invitrogen, Waltham, Massachusetts, USA) and the Qubit 2.0 fluorometer (Thermo Fisher Scientific, Waltham, Massachusetts, USA). PCR reactions were finally purified using AMPure bead purification protocol and samples with the concentration of 5 ng or more underwent the next steps of PacBio sequencing (PacBio Biosciences Inc, California, USA). SMRTbell libraries were prepared from the amplified DNA by blunt-ligation according to the manufacturer’s instructions. Circular consensus sequence (CCS) reads were generated from the raw PacBio sequencing data using the standard software tools provided by the manufacturer.

### Functional profiling using mRNA sequencing protocol

Eukaryotic cytoplasmic and mitochondrial ribosomal RNA as well as bacterial ribosomal RNA was depleted with removal probes and magnetic beads using Ribo-Zero Kit Epidemiology (Illumina, San Diego, CA, USA). The quality and quantity of total RNA, enriched mRNA and cDNA was assessed using the 2100 Bioanalyzer instrument and dedicated kits: RNA 6000 Pico and High Sensitivity DNA (Agilent Technologies Inc, Santa Clara, USA). Libraries for transcriptomics were generated using ScriptSeq v2 RNA-Seq (Illumina). Libraries were sequenced in single-end mode (50 or 68 base pairs) on an Illumina HiSeq 2500 sequencer using the TruSeq SBS Kit v3—HS (Illumina).

### 16S rRNA gene and RNA sequence reads pre-processing and analysis

CCS sequences from PacBio sequencing were analyzed using an in-house pipeline^58^. Sequences were identified to species level taxa by analysis of BLAST results obtained from a modified bacteria-only version of the SILVA SSU Ref_NR 99 database^59^ enriched with Human Oral Microbiome Database (HOMD)_16S_rRNA RefSeq Version 14.51 sequences^60^, and the All-Species Living Tree Project (LTP) database version LTPs128_SSU^61^, supplemented with unnamed and phylotype sequences from HOMD 16S_rRNA RefSeq. Reads that could not be reliably assigned to one species were clustered into 97% identity OTUs using UPARSE^62^ as implemented in USEARCH 10.0.240. Higher taxonomic levels were assigned using the RDP classifier version 2.13 at a cutoff bootstrap confidence of 80%^63^. Reads that were identified as typical contaminants using blanks and correlation analysis^64^ were removed.

BCL files were converted to FASTQ files (single-end reads) using bcl2fastq conversion software version v2.20.0.422 (Illumina). Sequencing read quality control was performed with fastp (ultra-fast all-in-one FASTQ preprocessor^65^). Reads of human origin were removed by a mapping approach incorporating bwa-samse (bwa-0.7.17, build 0.7.17-r1188^66^) against the human reference genome sequence GRCh38.p13 (GCA_000001405.28, NCBI RefSeq, PRJNA31257 “The Human Genome Project, currently maintained by the Genome Reference Consortium”).

To create a human oral-cavity-specific metagenome reference set, sequences from two publicly available datasets were selected and merged to form a combined database. The first source was the expanded Human Oral Microbiome Database (eHOMD, Rev: 2021-10-01, full eHOMD contig set), a widely known reference database in the field of oral metagenome analysis, which includes 2087 bacterial genomes from the human mouth and aerodigestive tract^67^. The second source was the study of Pasolli *et al.* 2019^68^, originally comprising 9,428 metagenomes and 154,723 metagenome-assembled genomes (MAGs)grouped into species-level genome bins (SGBs). From this source, only the 7897 MAGs found in the study’s oral-cavity-derived samples were selected. The sequences from both sources were combined into a combined oral metagenome reference set, grouped into 95% average nucleotide identity (ANI) taxonomic units (TUs) representing the species level, and annotated with the Prokka pipeline^69^. To extend the annotation and to assign additional EC numbers for further analysis, the eggnog annotated orthology database was applied^70^. The original taxonomy annotation of the eHOMD source combined with Prokka and eggNOG annotation form a comprehensive informative basis for the reference database created (taxonomy and functional annotation). Finally, in order to exclude taxa not present in our patient cohort, the TUs of the combined human oral metagenome reference set were selected if they were sufficiently covered by reads from our oral metagenome sample stock using a mapping approach (BWA-samse^67,71^). For the final mapping of the sample reads, all rRNA-coding genes were filtered out based on the respective annotation.

In order to determine read counts for the transcriptomic analysis sample reads were mapped to the protein-coding gene sequences from the selected TUs (SGBs) of the customized human oral reference set using BWA-samse. To avoid a biased counting for the transcript abundance of features, the reads mapped to multiple features were randomly distributed between them during the mapping of the sample reads against the customized human oral reference. Read counts were determined from the aligned sequencing read files applying htseq-count program from the HTseq package^72^. In order to reduce complexity and focus the feature definition on a functional level, the counts were summed up according to their gene-EC relationship. Genes that were annotated with multiple EC numbers were counted at the EC hierarchy levels at which the numbers were still identical. If the EC numbers of one gene still differed in the second-highest hierarchical level, the read counts for this gene were assigned to the “Unknown” group.

### Statistical analysis

Multivariate analysis of the species, class and genus level taxonomic data and EC based functional data was performed using ‘PRIMER 7 with PERMANOVA+ add-on’ software^73,74^. Principal co-ordinate analysis (PCoA) was performed for visualization and exploration of the Bray Curtis dissimilarities calculated on log(x+1) transformed data. A Permutational Multivariate Analysis of Variance (PERMANOVA)^75^ test was conducted using 99999 permutations to test the differences among groups. Similarly, a multivariate extension of Levene’s test called ‘PERMDISP’^76^ was used to test the with-in group dispersion of homogeneity. The group differences based on diagnosis groups were demonstrated using the PERMANOVA+ submodule canonical analysis of principal coordinates (CAP) on Bray Curtis dissimilarity matrices of log(x+1) transformed data of microbial abundances^77^. We used the Kyoto Encyclopedia for Genes and Genomes (KEGG) pathway mapper^78^ and iPath 3.0 (https://pathways.embl.de/)^79^ to integrate the ECs from our set onto the metabolic pathway map in order to identify and explore the pathways that were associated with healthy and peri-implantitis samples. Briefly, we characterized pathways that included ECs with a strong correlation with CAP axis (r > 0.4) separating the diagnosis groups and those with significant difference in their relative abundances in health and peri-implantitis (Mann-Whitney U test, *P*<0.05). Then, we used the https://www.kegg.jp/kegg/mapper/search.html in Reference mode to obtain in-depth information about different metabolic pathways. Visualization of pathways was created using iPath 3.0. For univariate analyses, we routinely applied Mann-Whitney U test. The adjusted p-values were calculated using the Benjamini-Hochberg correction.

### Curated amino acid metabolic pathway database reconstruction

Manual curation of in-silico amino acid metabolic pathways was systematically performed by employing 1. literature search for known amino acid metabolism by oral bacteria^21,27,80,81,82,83^, 2. automated web-based tool GapMind^84^ for annotating amino acid metabolism in bacteria^85,86^ and 3. MetaCyc^87^, which is a highly curated, non-redundant reference database of small-molecule metabolism. Unlike functional analysis of metatranscriptomes, this analysis is based on a reconstruction of metabolic pathways in individual genomes of microbes within biofilms and further prediction of possible interactions between different taxa. Biofilm bacterial requirements for 10 amino acids were predicted for (n = 20) bacterial classes. For each amino acid, we performed manual refinement to reconstruct the most plausible and complete (expressing all the ECs between substrate and end-product) metabolic pathways in bacteria. Some amino acids (glutamate, aspartate and alanine) could not be analyzed as they are not annotated in GapMind due to challenges with the annotation of related transaminases^84^. We focused on amino acids which related pathways were highly expressed in our dataset. Both anabolic and catabolic pathways for each amino acid and their transcribing taxa were aggregated into a count matrix table **(Extended data H)** that included the expression levels of ECs in all samples. To cross-validate our curated database of amino acid metabolism across bacterial taxa, we ran GapMind tool for the 3 genomes each from the representative bacterial species belonging to the most highly abundant genera from our RNAseq data (**Supplementary Fig. 7**). To infer the amino acid ecology, we calculated the folds between specific relative activity (related to either anabolism or catabolism of specific amino acids) for a given taxon and relative total activity of this taxon in biofilm. Low and high anabolism and high catabolism for specific amino acid-taxon combinations was defined if absolute log2 fold of specific to total activity was higher than 1.5. Relationships between amino acids and taxa were visualized as a network graph.

### Machine learning analysis

Linear and non-linear methods were used to build predictive models to identify biomarkers with high accuracies for diagnosis prediction (**Supplementary Fig. 1**). We employed CAP, which works as linear discriminant analysis using ‘PRIMER 7 with PERMANOVA+ add-on’ software, to identify the features with highest Spearman rank correlation to CAP1 axis, as potential candidates for biomarkers. We then built random forest (RF)^88^ models in R that were trained on our reduced set of full-16S and metatranscriptomic EC data. The goodness of fit of the models was assessed by training the model in a 20 times repeated 10-fold cross-validations and evaluating the resulting predictions through the mean area under the receiver’s operating characteristic curve (AUC). Additionally, for selection of optimum number of predictive features, we also employed recursive feature elimination (RFE)^89^ in R using Caret package with 10 features eliminated at each iteration. Hyperparameters were automatically tuned using a random grid search. The effect size and statistical significance of top biomarkers was calculated using Cohen d statistics and FDR corrected *p* values.

## Data availability

The raw data and the extended data files supporting the findings of this study will be made available by the corresponding author upon request. Full-16S and RNAseq data are deposited in the NCBI SRA as BioProject PRJNA1192962 and will be made available after publication of this work.

## Acknowledgement

SPS, SH & MSti would like to thank Deutsche Forschungsgemeinschaft (DFG, German Research Foundation) for funding (SFB/TRR-298-SIIRI – Project-ID 426335750). This study was also funded by the DFG under Germany’s Excellence Strategy - EXC 2155 - project number 390874280 (SH & MSti). We would like to thank Marly Dalton and Rainer Schreeb for technical assistance.

## Contributions

SH and MSti conceived and planned the study. PSD and JG collected the samples. AJ, SPS, TQ and XX performed 16S rRNA amplicons and RNA sequencing. AJ, SPS, MSte, IY and WB performed the bioinformatic analysis. Data interpretation and visualizations were performed by AJ and SPS. AJ and SPS wrote the manuscript and MSti revised it. All authors read and approved the final manuscript. AJ and SPS have contributed equally to the work.

