## Supplementary for "Integrative microbiome- and metatranscriptome-based analyses reveal diagnostic biomarkers for peri-implantitis"

Short title: Biomarkers for peri-implantitis

Amruta Joshi[1,2,*], Szymon P. Szafrański[1,2,3,*], Matthias Steglich[1,2,], Ines Yang[1,2], Taoran Qu[1,2], Paula Schaefer-Dreyer[1], Xing Xiao[1,2], Wiebke Behrens[1,2], Jasmin Grischke[1], Susanne Häussler[3,4,5,6], and Meike Stiesch[1,2,3,$]

[1]Department of Prosthetic Dentistry and Biomedical Materials Science, Hannover Medical School, Hannover, Germany

[2]Lower Saxony Centre for Biomedical Engineering, Implant Research and Development (NIFE), Hannover, Germany

[3]Cluster of Excellence RESIST (EXC 2155), Hannover Medical School, Hannover, Germany

[4]Department of Molecular Bacteriology, Helmholtz Centre for Infection Research, Braunschweig, Germany

[5]Institute for Molecular Bacteriology, Twincore, Centre for Clinical and Experimental Infection Research, Hannover, Germany.

[6]Department of Clinical Microbiology, Copenhagen University Hospital - Rigshospitalet, Copenhagen, Denmark

[*]Amruta Joshi and Szymon P. Szafrański contributed equally

[$]correspondence to: Prof. Dr. Meike Stiesch, Department of Prosthetic Dentistry and Biomedical Materials Science, Hannover Medical School Carl-Neuberg-Str.1 30625 Hannover, Germany;

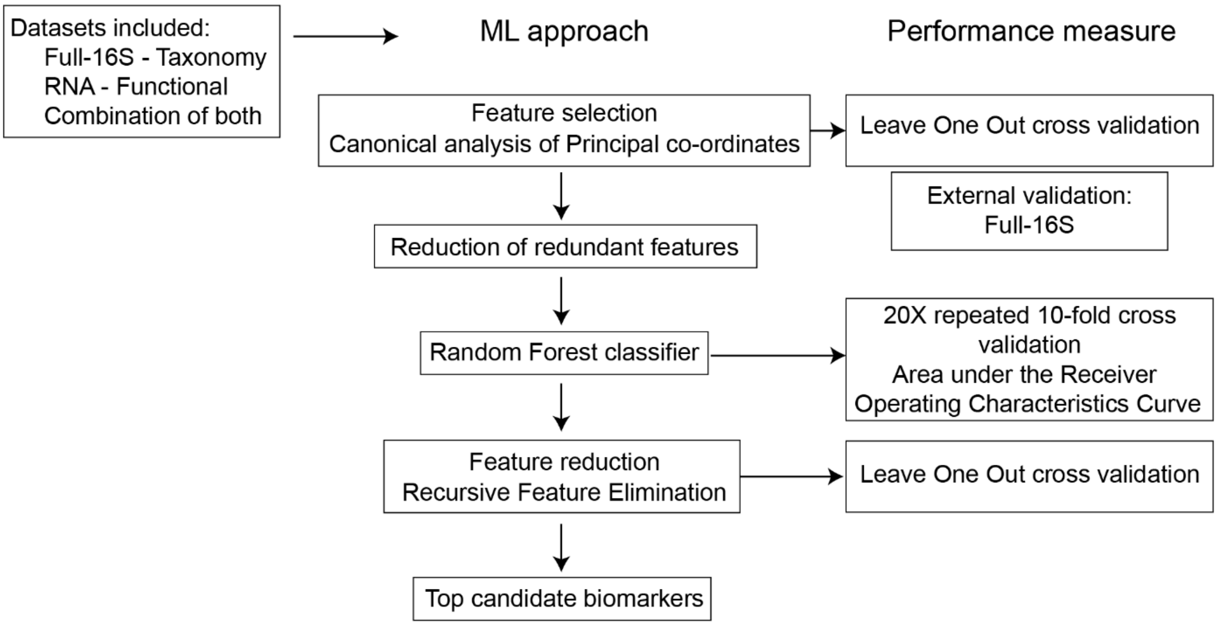


**Supplementary Fig. 1 Machine learning workflow for biomarker identification and validation.**

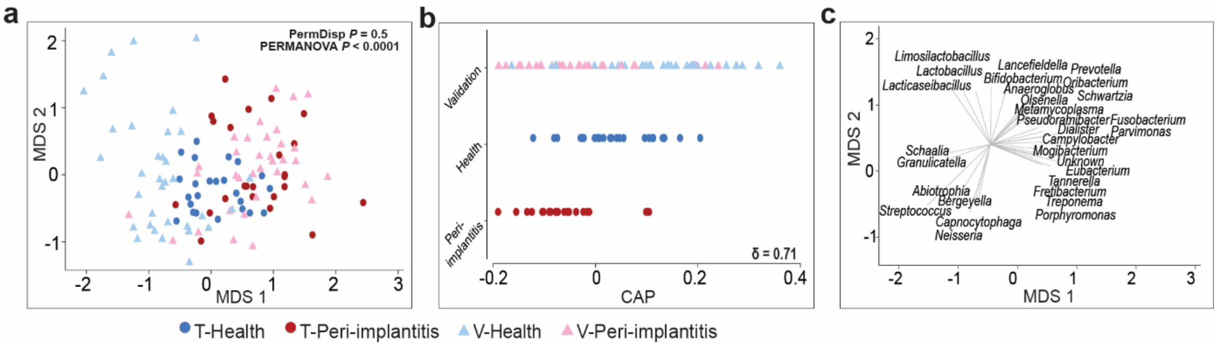


**Supplementary Fig 2. Separation between diagnosis groups based on full-16S genus-level taxa.**

**a** Non-metric multi-dimensional scaling (nMDS) plot based on Bray-Curtis dissimilarity matrices of 16S rRNA gene amplicon data, aggregated to genus level features of the training and validation set samples. **b** Canonical analysis of principal co-ordinates (CAP) constrained ordination plot of genus level, illustrating that CAP axis that maximizes the separation of the two 'a priori' groups based on diagnosis. New observations (validation samples) used to test the performance of CAP model shows the successful allocation of validation samples to their respective diagnosis groups. **c** Vector overlays on nMDS shows the genera with highest correlation to the CAP1 axis that best separates the diagnosis groups as seen in (b).

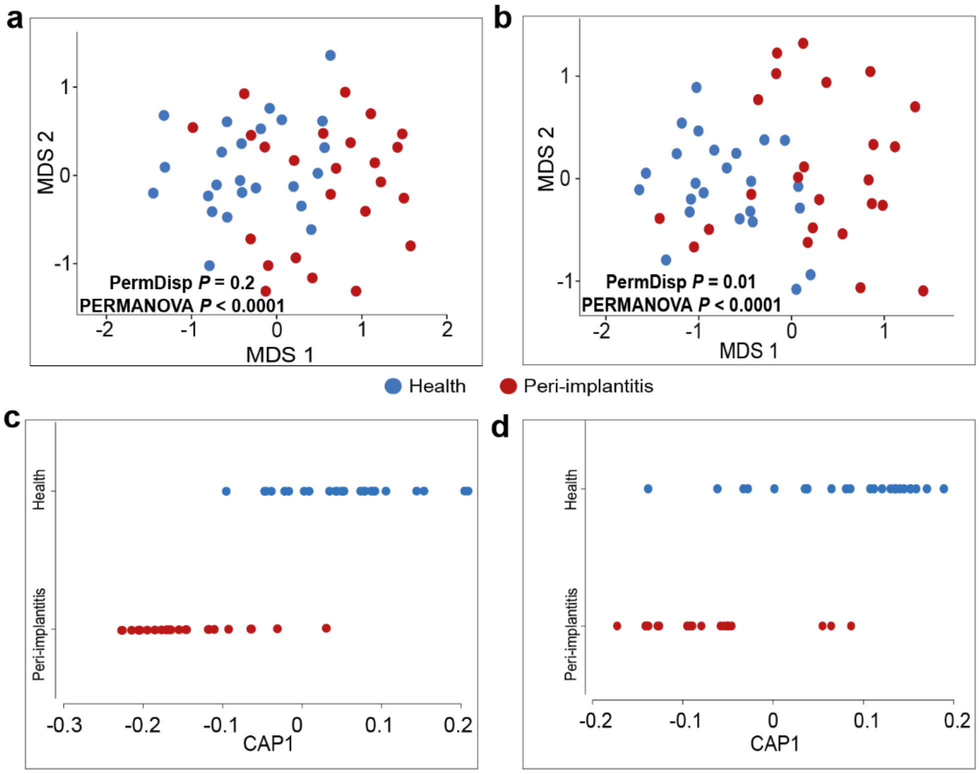


**Supplementary Fig 3. Separation between diagnosis groups based on combined datasets.**

Non-metric multi-dimensional scaling (nMDS) plots **(a)** species + EC4 combination and **(b)** genus + EC4 combination. Canonical analysis of principal co-ordinates (CAP) plots of **(c)** species + EC4 combination and **(d)** genus + EC4 combination, illustrating the CAP1 axis that maximizes the separation of the two 'a priori' groups based on diagnosis.

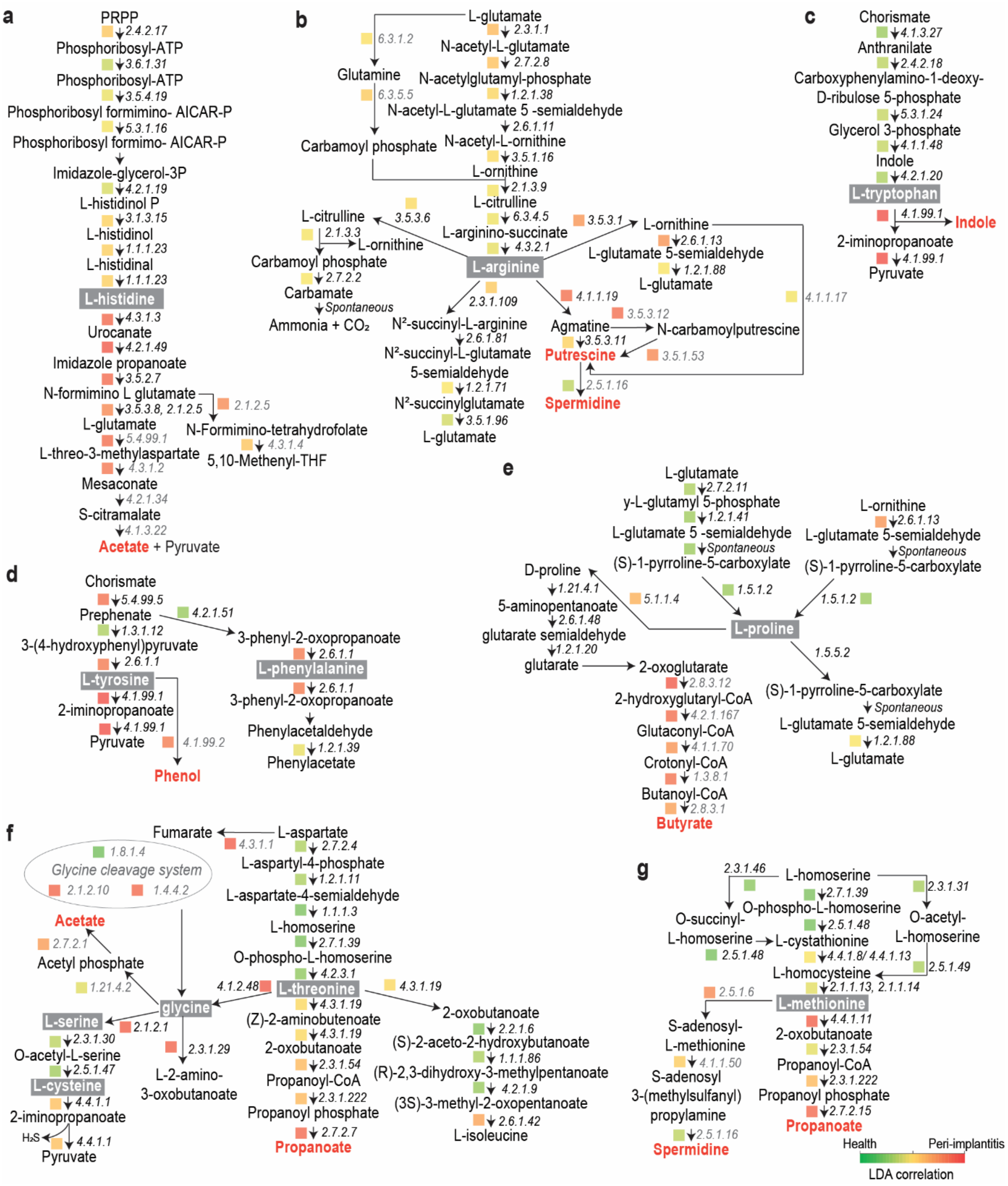


**Supplementary Fig. 4. High-resolution view of the curated pathways for selected amino acids based on RNAseq ECs.** Anabolic and utilization pathways for **a.** histidine, **b.** arginine, **c.** tryptophan, **d.** tyrosine and phenylalanine, **e.** proline, **f.** threonine, glycine, serine and cysteine, and **g.** methionine were identified using metatranscriptomic data. Colour scale indicates the ‘canonical analysis of principal co-ordinates (CAP)’-based linear discriminant analysis (LDA) correlation of each EC to the CAP1 axis that best separates the diagnosis groups.

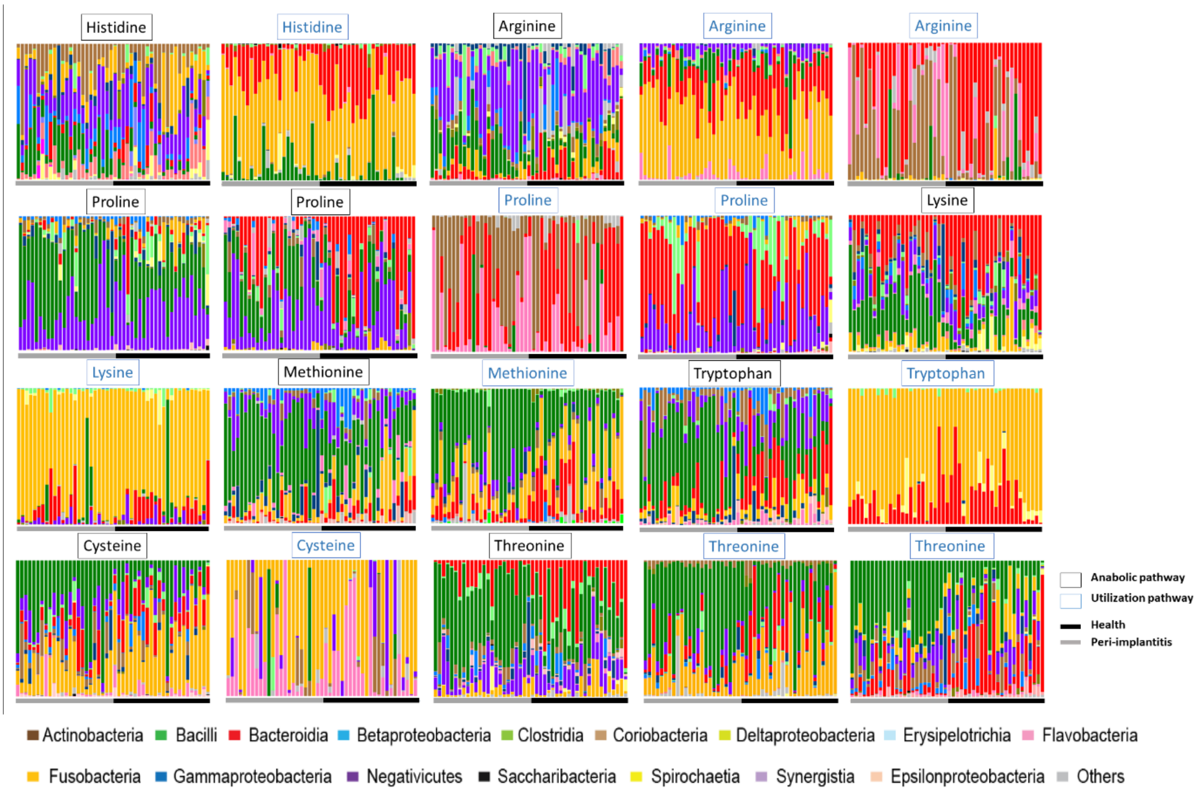


**Supplementary Fig. 5. Relative abundances of bacterial taxonomic classes contributing to amino acid anabolic and utilization activities in each sample from healthy and peri-implantitis groups.** Mean relative abundances of the contributing bacterial classes to selected pathways are shown in Fig. 6 of the manuscript.

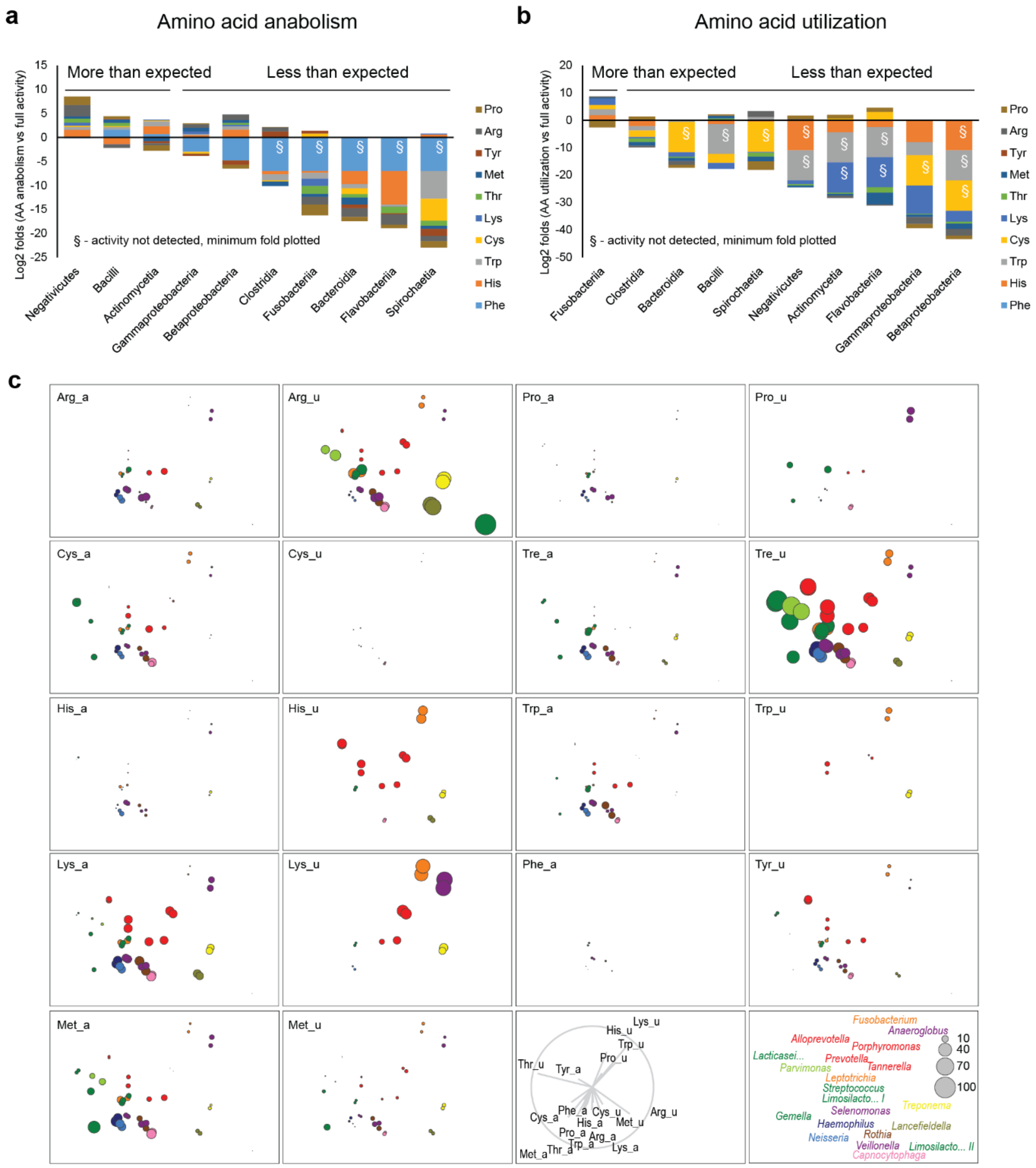


**Supplementary Fig. 6. Taxon-specific strategies for amino acids metabolism.**

**a** Log 2-fold change plotted for transcriptional anabolic activities of specific bacterial classes against their full amino acid activity. **b** Log 2-fold change plotted for transcriptional catabolic activities of specific bacterial classes against their full amino acid activity. **c** Non-metric multi-dimensional scaling (nMDS) plots depicting amino acid metabolism for 22 genera for specific anabolic and utilization pathways. Vector overlays for pathways and genera plotted at the end. Bubble sizes reflect % of reads (EC-based) assigned to specific amino acid-related activities. Genera were split by health status (see **Fig. 7b**).

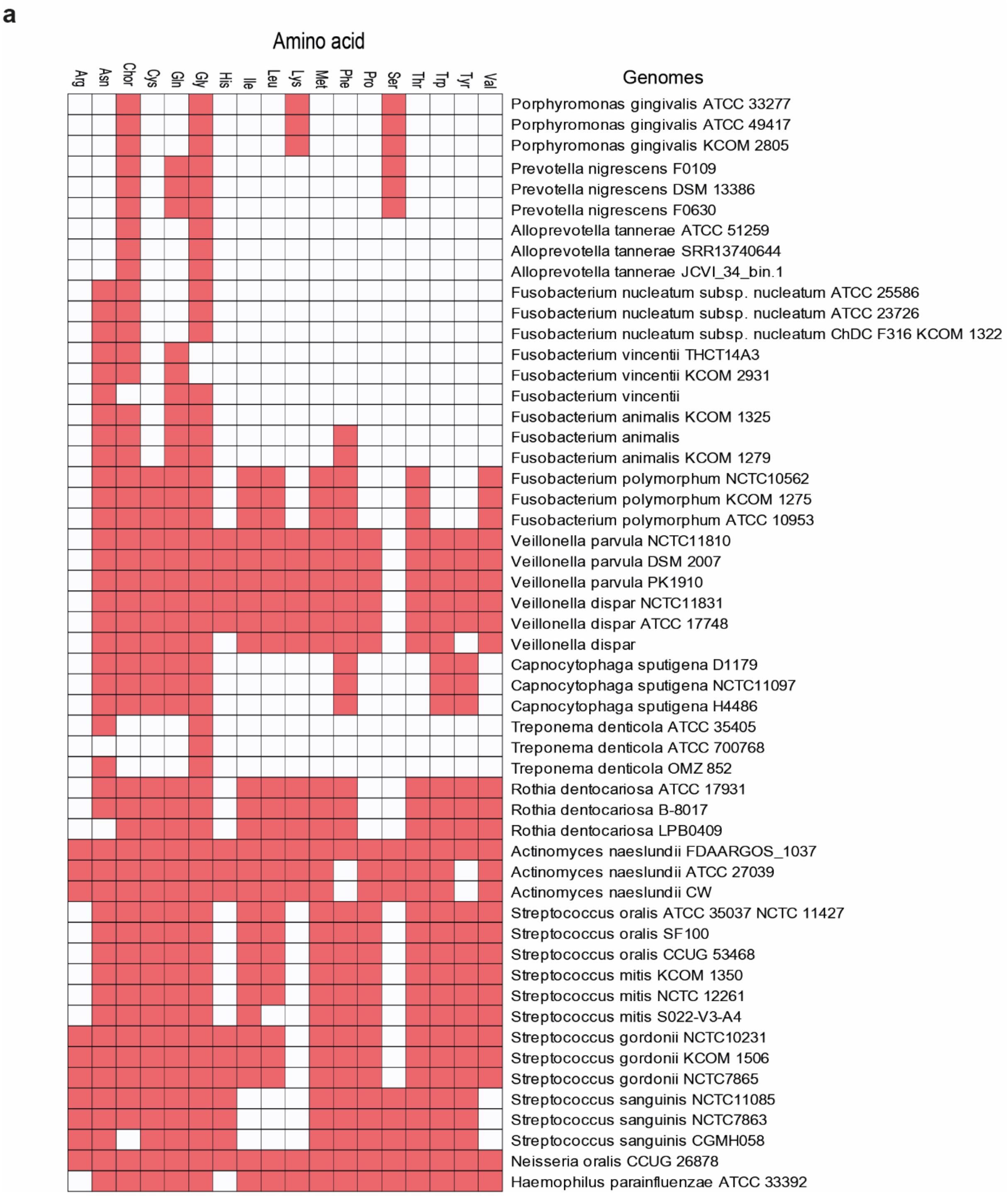


**Supplementary Fig. 7. GapMind based cross-validation of the curated amino acid biosynthetic activities of the representative species from the 20 most highly abundant genera detected in our study.**

### Extended Data

**A. Demographic and clinical characteristics of the study cohort.**

**B. Demographic and clinical characteristics of the validation cohort.**

**C. Full-16S based species level composition.** Statistical significance, CAP-based LDA correlation, species-level mean abundances in healthy and peri-implantitis groups

**D. Full-16S based genus level composition.** Statistical significance, genus-level mean abundances in healthy and peri-implantitis groups

**E. Full-16S based class level composition.** Statistical significance, class-level mean abundances in healthy and peri-implantitis groups.

**F. Metatranscriptome-based features (Enzyme Commission numbers - EC).** Statistical significance, CAP-based LDA correlation, Mean 4-digit level EC abundances in healthy and peri-implantitis groups.

**G. Metatranscriptome-based class level taxonomic features.** Statistical significance, class level mean abundances in healthy and peri-implantitis groups.

**H. Curated amino acid metabolism dataset.** Mean abundances of ECs aggregated to anabolic and catabolic pathways and their transcribing active taxa.
